## Supplementary figures and images for "A public resource of single cell transcriptomes and multiscale networks from persons with and without Alzheimer’s disease"

### Supplementary Figure 1

#### Supplementary Figure. 1

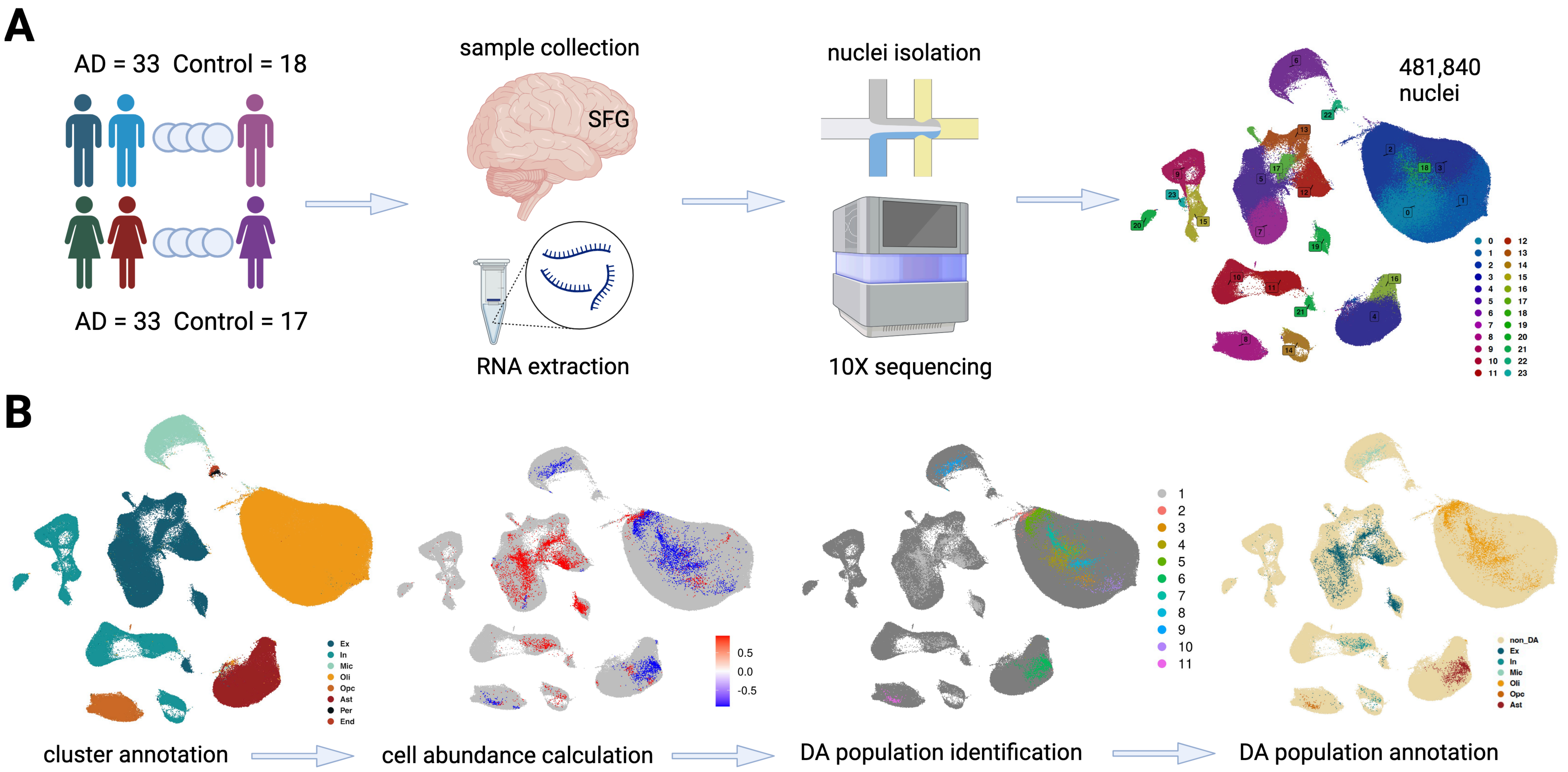

### Supplementary Figure 2

#### Supplementary Figure. 2

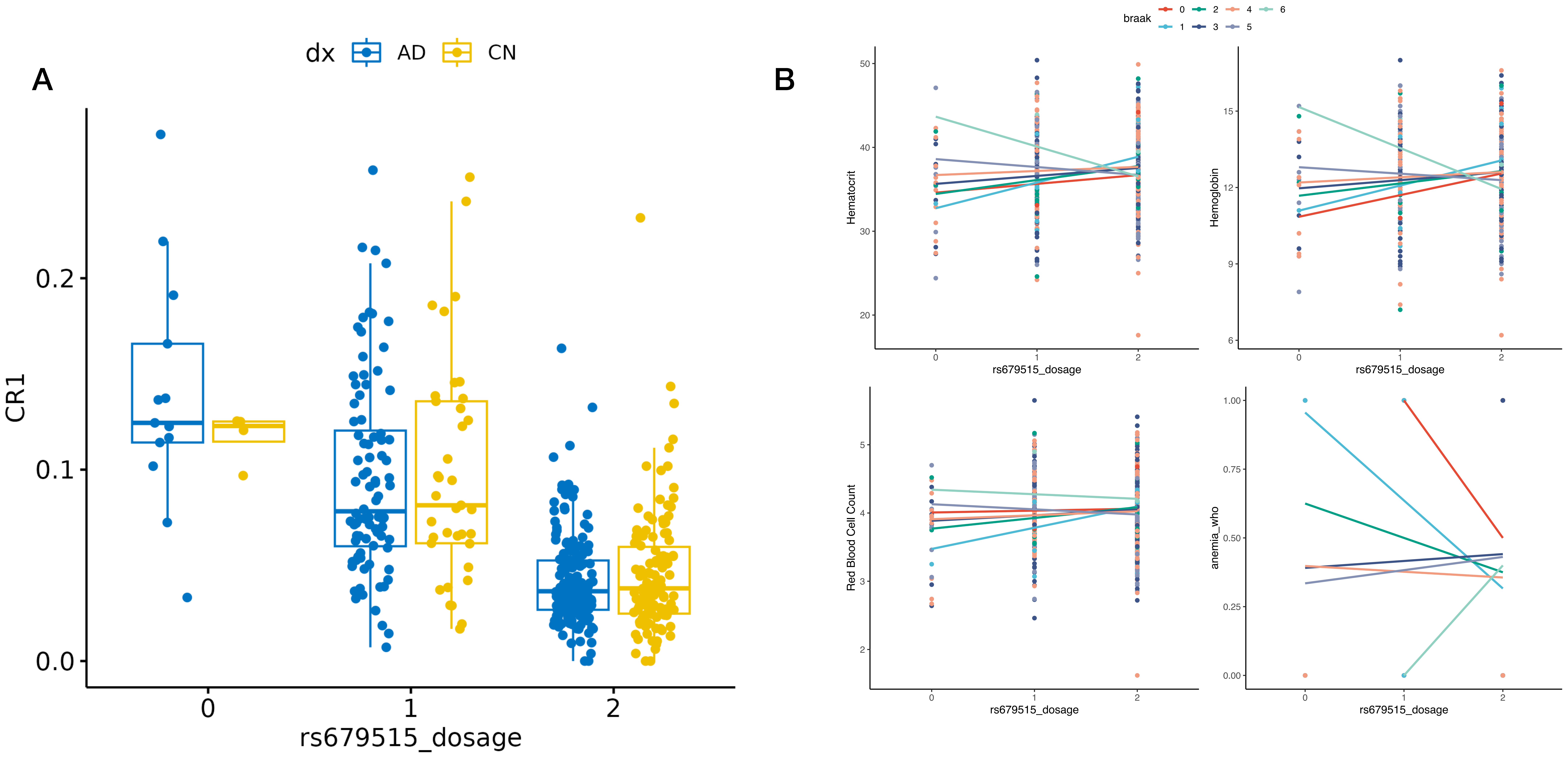

### Supplementary Figure 3

#### Supplementary Figure. 3

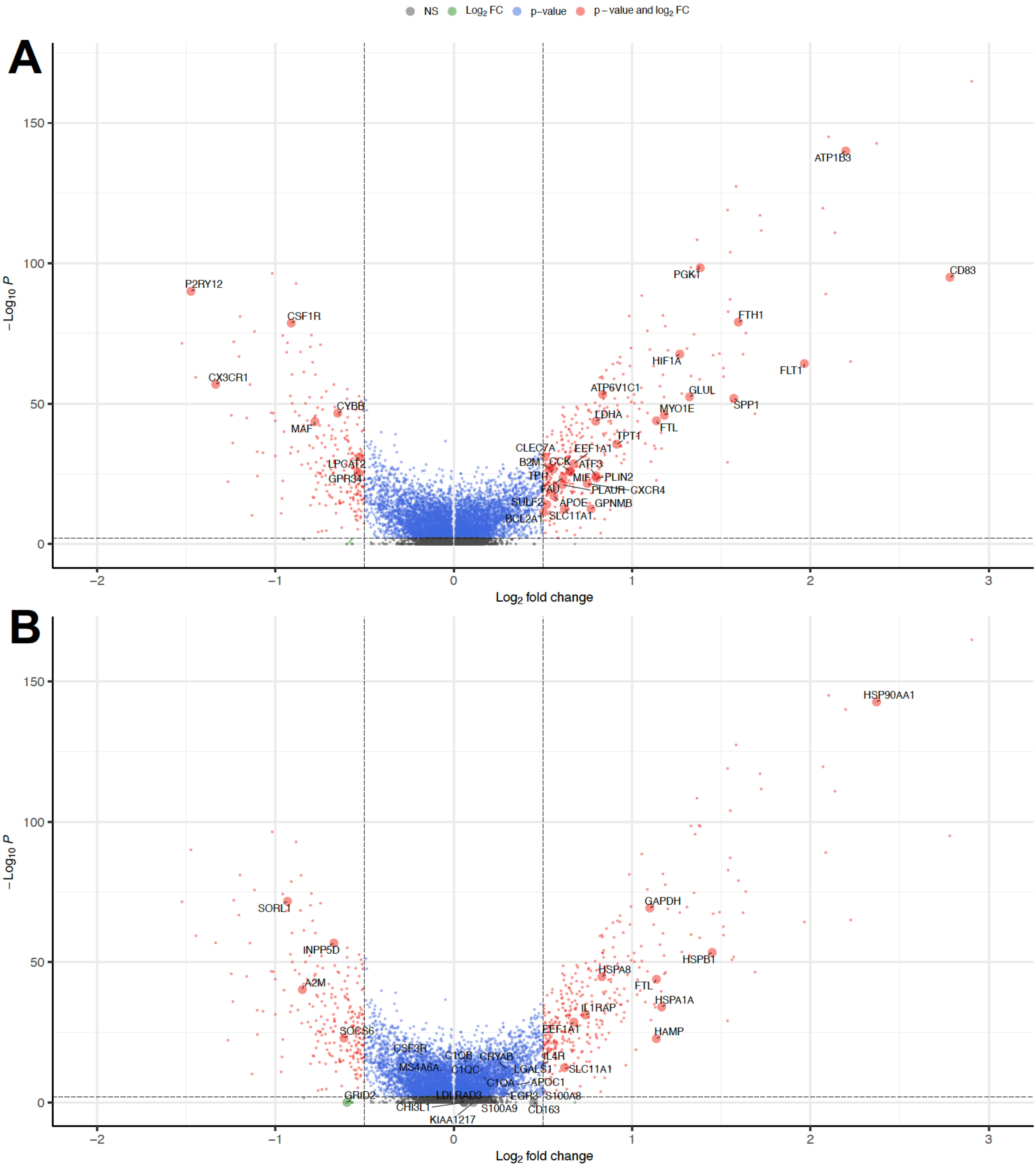

### Supplementary Figure 4

#### Supplementary Figure. 4

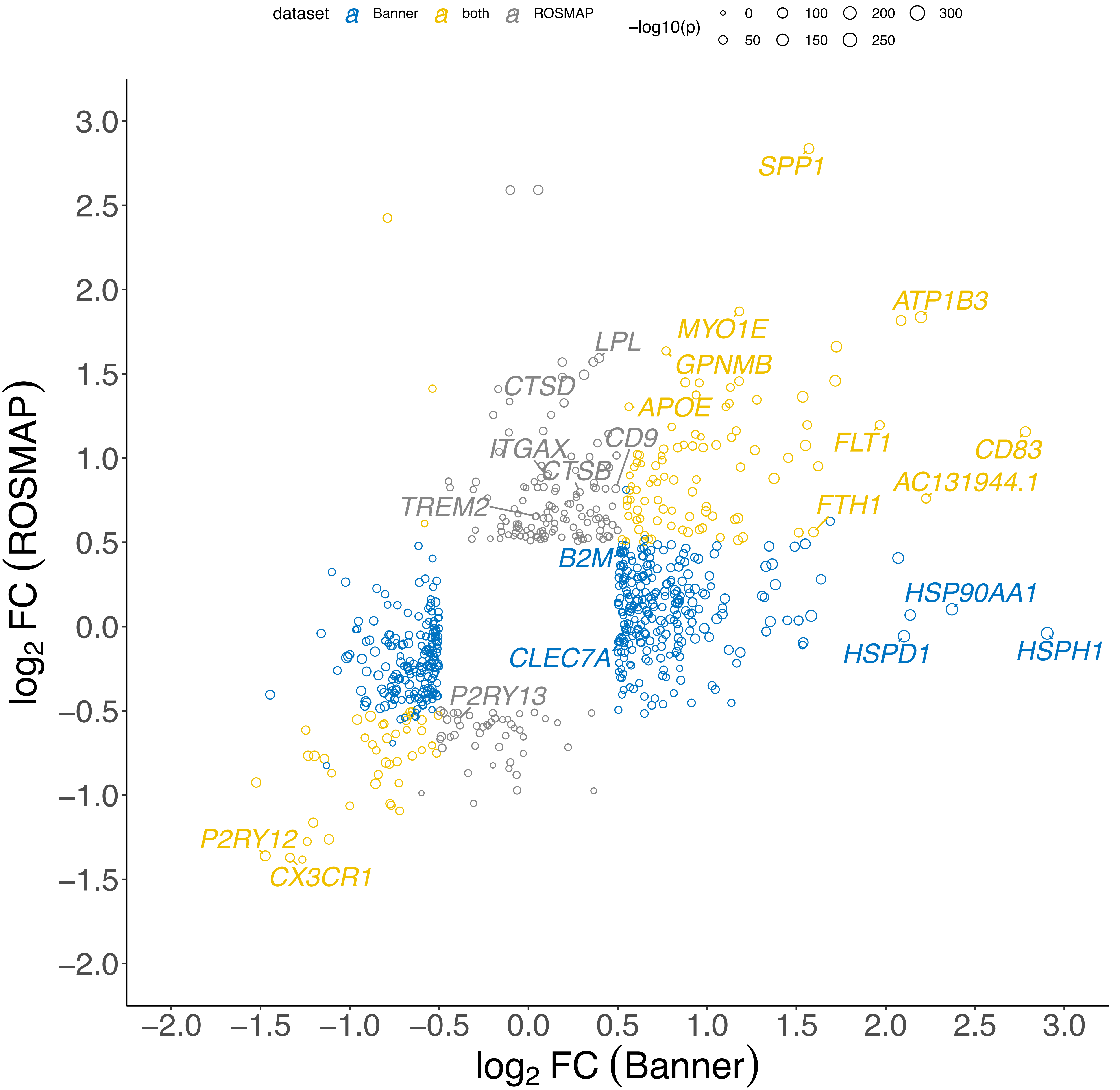

### Supplementary Figure 5

#### Supplementary Figure. 5

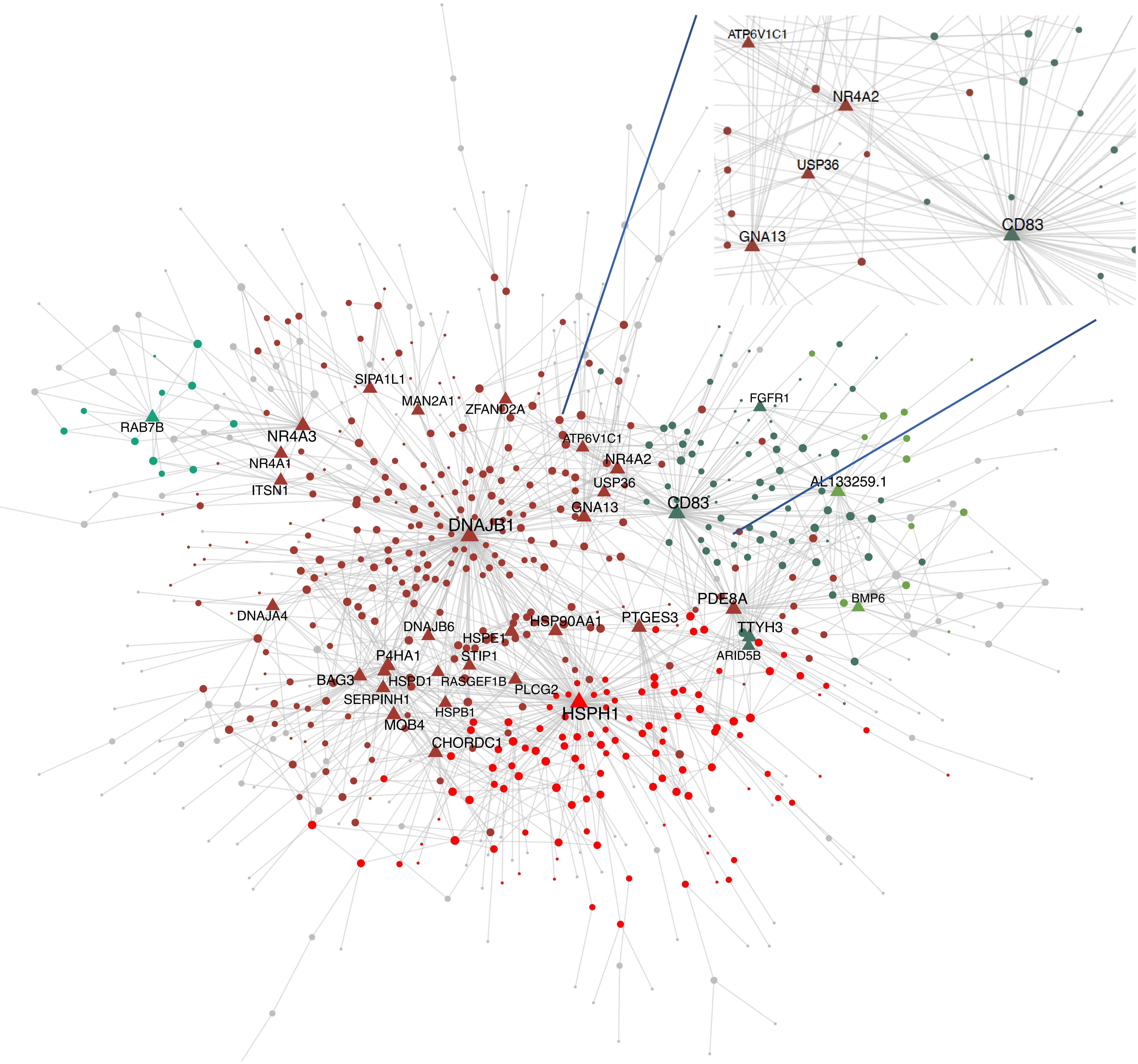
